# Properties of Predictive Gain Modulation in a Dragonfly Visual Neuron

**DOI:** 10.1101/496885

**Authors:** J. M. Fabian, J. R. Dunbier, D. C. O’Carroll, S. D. Wiederman

**Author notes:** Corresponding Author: Joseph Fabian.

## Abstract

Dragonflies pursue and capture tiny prey and conspecifics with extremely high success rates. These moving targets represent a small visual signal on the retina and successful chases require accurate detection and amplification by downstream neuronal circuits. This amplification has been observed in a population of neurons called Small Target Motion Detectors (STMDs), through a mechanism we termed predictive gain modulation. As targets drift through the receptive field responses build slowly over time. This gain is modulated across the receptive field, enhancing sensitivity just ahead of the target’s path, with suppression of activity elsewhere in the surround. Whilst some properties of this mechanism have been described, it is not yet known which stimulus parameters are required to generate this gain modulation. Previous work suggested that the strength of gain enhancement was predominantly determined by the duration of the target’s prior path. Here we show that the predictive gain modulation is more than a sluggish build-up of gain over time. Rather, gain is dependent on both past and present parameters of the stimulus. We also describe response variability as a major challenge of target detecting neurons and propose that the predictive gain modulation’s role is to drive neurons into response saturation, thus minimising neuronal variability despite noisy visual input signals.

## Introduction

Dragonflies detect and capture small moving prey, and pursue conspecifics for mating and territorial defence. During predatory flights, the dragonfly’s target rarely spans more than 1° of visual space, thus stimulating only two or three ommatidia of the compound eye (Lin and Leonardo, 2017). A population of target-detecting neurons, which we refer to as Small Target Motion Detectors (STMDs), have been identified in the optic lobe of several inspect species, including the dragonfly (O’Carroll 1993; Geurten et al. 2007). These STMDs are responsive to target contrasts as low as 1.3% (Wiederman, Fabian et al, 2017), matching the sensitivity of fly ‘optic-flow’ neurons that are responsive to large moving patterns (Harris et al. 2000). However, such responses to large objects are the result of many presynaptic units being integrated over space, each viewing different portions of the same feature (Bausenwein et al. 1992). STMDs do not have a similar advantage, as each unit encounters a small and noisy signal which cannot be pooled. It is at an individual photoreceptor’s signal detection limit where target sizes are able to elicit STMD responses and generate pursuit behaviour (Rigosi et al. 2017).

We recently described a predictive mechanism employed by STMDs that increase responses to these low signals. Following target onset, spiking activity is initially weak, building in strength over several hundred milliseconds when the target’s trajectory is continuous in space and time (Nordström et al. 2011; Dunbier et al. 2012). This property, we termed ‘facilitation’, is matched to a large improvement in contrast gain, with detection thresholds improved 5-fold following the longer target motion (Wiederman, Fabian et al. 2017). The facilitation results in a local gain increase situated slightly ahead of a target’s current position, and a global suppression in the extended surround. This ‘predictive gain modulation’ spreads forward in space following an occlusion, matched to the target’s direction of travel. As a result, signal strength is improved for more realistic, continuous target trajectories, whilst background distracters and false positives are suppressed.

Psychophysical experiments reveal that humans use information of prior trajectory to improve their ability to detect and track moving targets (Watamaniuk et al. 1995; Watamaniuk and McKee 1995). For example, during target occlusions human account for target attributes, such as velocity, in their predictions (Bennet and Barnes 2003). Similarly, observers looking at a scene use specific feature attributes such as their size, direction of movement and colour, to ignore distracters and enhance the saliency of a target of interest (Saenz et al, 2002).

For the dragonfly, the angular size and velocity of a moving target may change dramatically throughout a pursuit. Therefore, robust interception of the target’s location should be resilient to a wide range of stimulus properties. Here we describe attributes of the target and its trajectory that drive the visual neuron’s predictive gain modulation. These results provide insight into how neuronal processing underlies fundamental behavioural tasks, such as predicting the future location of a moving target in the visual environment.

## Materials and Methods

### Animals and Electrophysiology

These data are from intracellular recordings in a total of 31 male wild caught adult *Hemicordulia*, gathered from the Adelaide Botanic Gardens. Animals were immobilized with a 1:1 bees wax-rosin mixture (Colophony Kolophonium, Fluka Analytical), and fixed to an articulating magnetic stand. The animals head was tilted forward to expose the posterior surface of the head capsule, before dissecting a small hole in the cuticle directly over the left lobula complex.

We used a Sutter Instruments P-97 electrode puller to taper Aluminosilicate electrodes, which were backfilled with 2M KCl solution (Potassium chloride, AnalaR). We step electrodes through the neural sheath and into the proximal lobula complex with a piezoelectric stepper (Marzhauser-Wetzlar PM-10). Electrodes had a typical electrical resistance of between 50-150 MΩ.

Freshly penetrated neurons were first presented a series of visual stimuli for neuronal classification, including small targets, bars, gratings and patterns. We identify CSTMD1 by its small target selectivity, large characteristic receptive field and action potential waveform (Geurten et al., 2007). Intracellular responses were digitized at 5 kHz with a 16-bit A/D converter (National Instruments) for off-line analysis with MATLAB.

### Visual Stimuli and experiment design

Visual stimuli were presented on high definition LCD monitors (144-165 Hz) placed 20 cm from the animal, centred on the animal’s frontal midline. Stimulus scripts were written using MATLAB’s Psychtoolbox and integrated into the data acquisition system. All stimuli consisted of 1.5° x 1.5° dark targets drifting vertically on a white background, with the exception of experiments presented in figure 4, where horizontally drifting targets of a series of heights were used. We used intermediate contrast (Weber contrast of 0.41) in order to prevent neuronal saturation masking the magnitude of the changes in gain. Experimental paradigms were randomised within each neuron and a minimum of 8 seconds rest was implemented to avoid habituation within and across trials. Data was only excluded due to pathological damage, as indicated by drifts in resting membrane potential, decreases in spike amplitude or excessively high spontaneous activity. Each experiment consisted of two stimulus types, probes and primers (Wiederman, Fabian et al., 2017). Probes are presented either alone or following a primer. Responses to ‘probe’ targets are weak when presented alone, and enhanced by varying amounts when preceded by a ‘primer’.

Spatiotemporal tuning of gain modulation: We replicated the experimental paradigm of Nordström et al (2011) by drifting targets on short (probe only) and long (primer plus probe) trajectories. The probe consisted of a 1.5°x1.5° target drifting at 40°/s for 200 ms, but unlike in previous work we used 16 different combinations of primer durations and distances, each terminating at the same position (n = 6 dragonflies).

Velocity gain map: We mapped the receptive field of CSTMD1 in one dimension using a series of 6 probes drifting vertically through the receptive field (n = 12 dragonflies). Each probe drifted at 45°/s for 135 ms, starting at different elevations separated by 6°, such that the end position of one probe was identical to the start position of the next. These probe trials were randomly interleaved with primed trials consisting of one of two primers – either drifting at 30°/s or at 60°/s (both with a duration of 500 ms). Primers terminated either 100 ms or 300 ms prior to presentation of the probe to test the spread of the predictive gain modulation.

Target primer height: To be consistent with prior work investigating target height tuning (Geurten et al., 2007; O’Carroll and Wiederman, 2014), we drifted targets horizontally. Target height tuning was measured by drifting probe targets (40°/s) at heights varied logarithmically (orthogonal to motion) between 0.16° and 20° (n = 15 dragonflies). We performed this height tuning at two target widths, 1.5° and 8°. Probes were presented alone, or following one of two priming targets with a size of either 1.5°x0.5° or 1.5°x5°(both drifting at 40°/s for 500 ms).

Neuronal Variability: Probes were moved vertically through the receptive field for 500 ms at 40°/s, either presented alone or preceded by 500 ms of primer motion (terminating at the probe’s start position). An individual dragonfly was presented both conditions of this experiment 40 times in randomised order, with each trial separated by 144 seconds to ensure that any neuronal variability observed was not due to local habituation. The same experiment was replicated 5-10 times in a further 8 dragonflies (n = 9 dragonflies total) to analyse the variability of gain modulation onset across different animals.

### Data analysis and statistics

Data presented in figures 1 to 4 are measured from the spiking activity within a 100 ms analysis window, beginning 50 ms following the onset of a probe. This analysis duration permits a spike rate calculation, whilst minimising the amount of ‘self-priming’ of the probe itself. Introducing a 50 ms offset accounts for CSTMD1’s latency (Nordström et al., 2011). Gain modulation is calculated by subtracting the response for probe-alone trials from their paired ‘primed’ probe response (corresponding time windows). This measure of change, allows us to compare gain modulation across animals, without confounding overall neuronal excitability. To compare height tuning between conditions, we display absolute responses to primed and unprimed probes.

**Figure 1:**
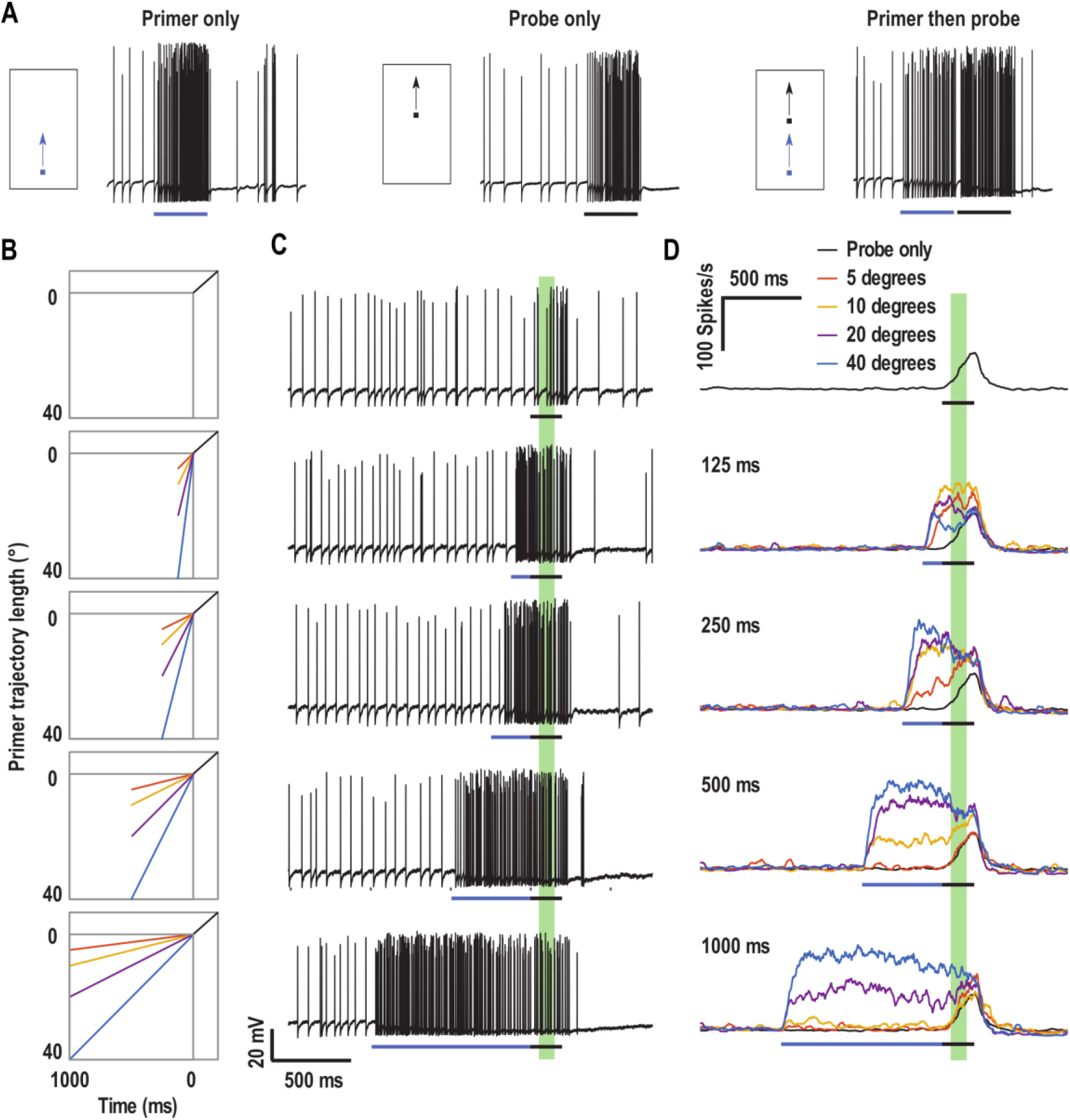
Priming targets facilitate the response to a probe. A) Probes drift on a short trajectory, preceded by either no stimulus or a priming target. The priming target generates predictive gain modulation, which can be observed by quantifying the change in response to the probe that follows. Blue and black stimulus bars represent primer and probe presentation respectively. B) The primer can be 1 of 16 different combinations of distances in space and durations in time. C) Example data traces from an individual CSTMD1 recording in response to varying primers. Spiking activity is calculated in a 100 ms window (green shaded area). D) The strength of the probe response varies dependent on the properties of the preceding primer (mean, n=6). Quantitate analysis of these data is presented in Figure 2.

**Figure 2:**
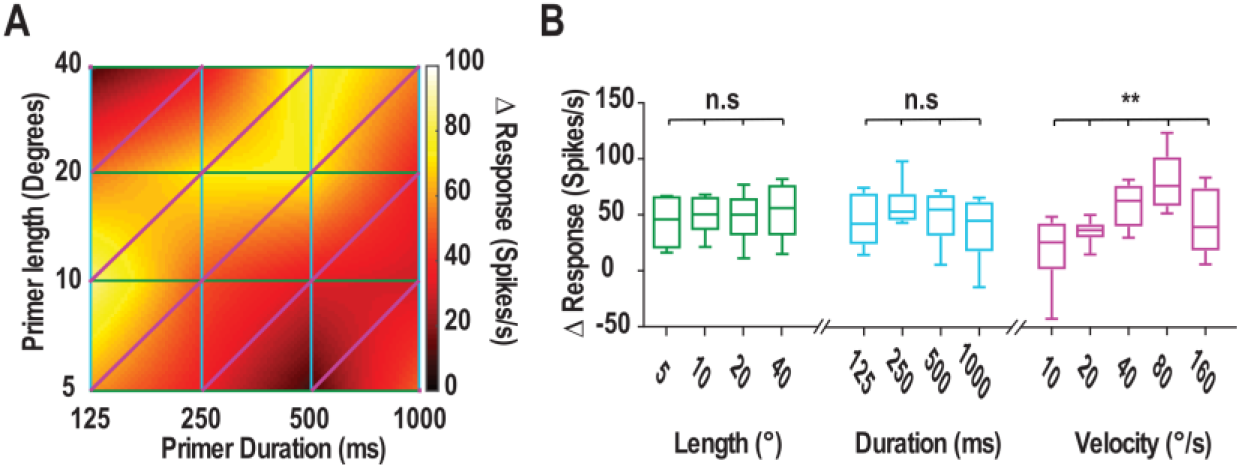
The relationship between primer length and duration on probe response gain. A) A 2D representation of the strength of probe gain modulation due to primers presented across different combinations of space and time. B) Each data point in (A) is arranged into groups of equal primer length, primer duration or primer velocity (n = 6 dragonflies). Duration and length alone do not have any clear effects on facilitation strength, but we observe significant variation based on priming target velocity.

**Figure 3:**
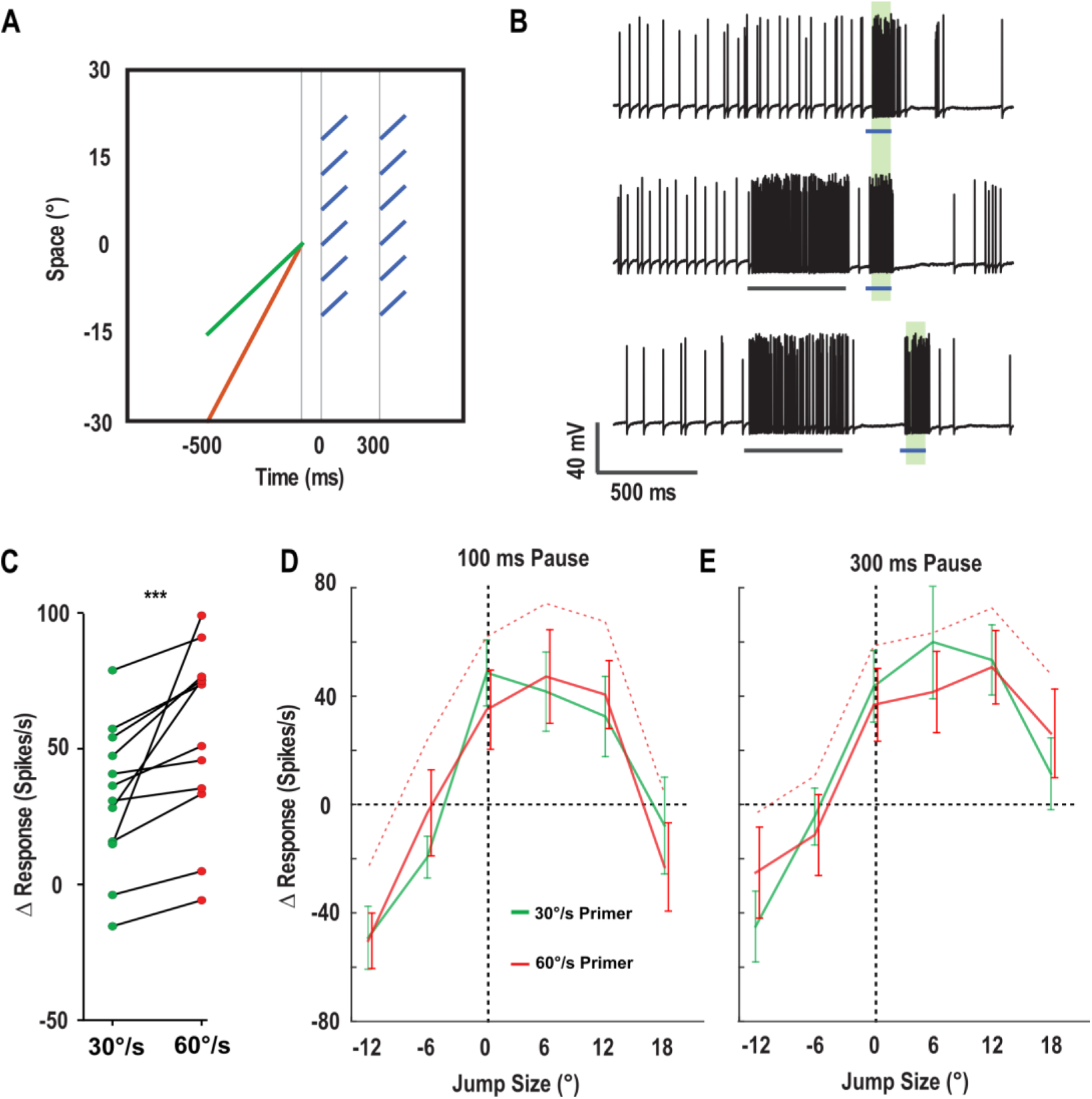
Primer velocity and the spreading focus of gain. A) The change in response to a series of probes (blue) in different locations is mapped following primers of two different velocities (30°/s green line; 60°/s red line), following either a 100 and 300 ms pause. B) Example responses to these stimuli from an individual CSTMD1 recording. Responses to probes are quantified in a 100 ms window, presented alone or following different primers. (C) Pooling the observed change in response for all forward probe trajectories (0-18°) reveals an increase in the strength of gain for probes presented following the faster primer (P < 0.001). This velocity dependent vertical offset (as observed in Figure 2) will confound our analysis of gain location. D) To account for this velocity dependency on overall gain (a vertical offset), we take the change induced by the faster primer velocity of 60°/s (red dashed line), and subtract the mean difference between 30°/s and 60°/s, thus creating the solid red line. This permits the plotting of the *location* of gain modulation at different ‘jump’ positions (0° is a continuous trajectory) after either a 100 ms pause or (E) at a 300 ms pause following the termination of the primer (mean ± SEM, n =12 dragonflies). This shows that although gain modulation spreads forward following the longer occlusion (300ms c.f. 100ms), there is no significant effect of primer velocity on the location of this spreading facilitation.

**Figure 4:**
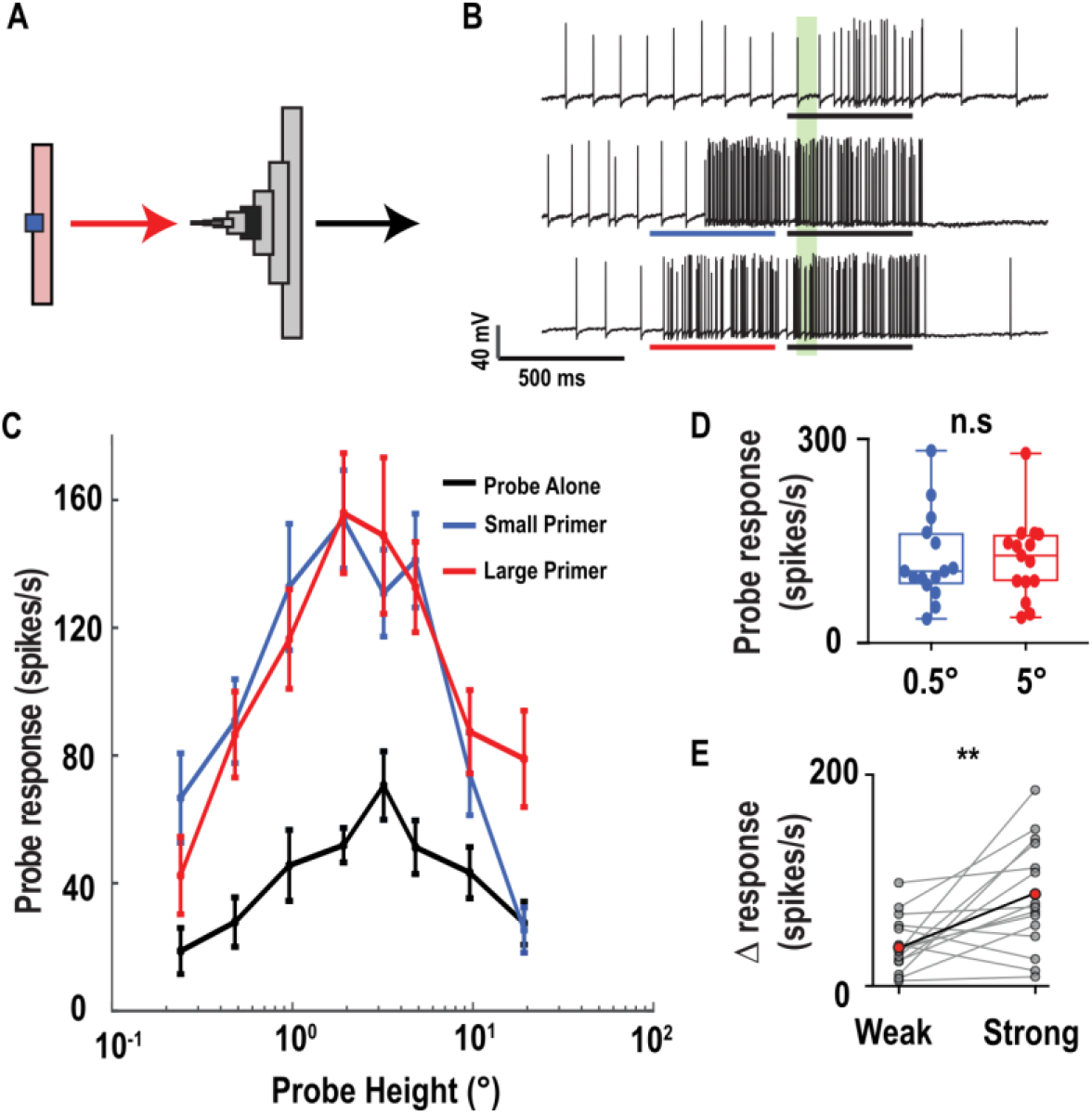
Target height tuning following priming with targets of different heights. A) Probe target height tuning is measured, either presented alone or primed by small (0.5°x1.5°) or large (5°x1.5°) targets. B) Example responses of an individual CSTMD1. Probe responses at the varying heights are quantified in a 100 ms window (green shaded region). C) Responses to probes of 8 varying heights, presented alone (black line) or following the two (small target, blue line; large target, red line) priming conditions (n = 15 dragonflies). The optimal height tuning is unchanged, irrespective of the primer size D) The response to both primers across all trials elicits similar responses, i.e. on either side of the height tuning curve. E) The change in response gain produced by identical primers for ‘weak’ probes (0.24°, 0.48°, 9.6° and 19.2° pooled) and ‘strong’ probes (0.96°, 1.92°, 3.2°, and 4.8° pooled). Grey points indicate individual trials, red points indicates mean.

In figure 5A-B we present results from an individual CSTMD1 recording, where we computed instantaneous spike rate for 40 trials of a 500 ms probe presented alone, as well as 40 trials of the same probe preceded by a 500 ms primer. Given the receptive field of CSTMD1 is inhomogeneous (Geurten et al., 2007), we normalised responses based on receptive field location. This is performed by dividing the instantaneous spike frequency of each individual short path trial by the mean instantaneous spike frequency for a primed probe presented at the same location. In figure 5C, the same experiment was performed in a total of 9 CSTMD1 recordings in subsequent animals, with the average probe onset presented for each cell. Figure 5E is generated by applying a 100 ms analysis bin, sliding in 1 ms increments. For each bin we compute the mean spike count across each trial, as well as the variance (σ^2^). These two values are also used to calculate the Fano Factor 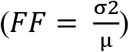.

**Figure 5:**
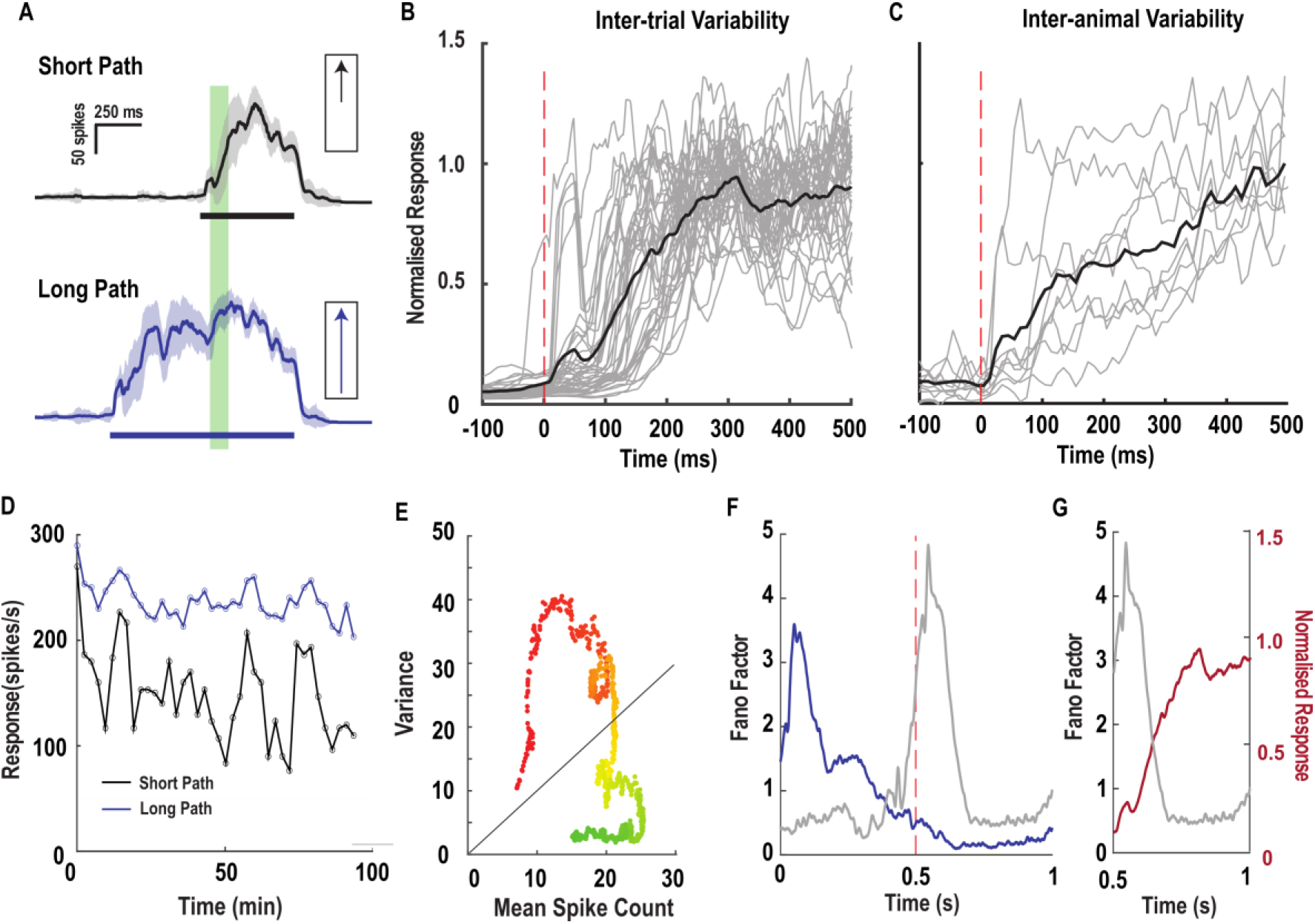
Neuronal variability and predictive gain modulation. A) Instantaneous spike frequency plots for the presentation of the short (top) and long (bottom) path target trajectories (shaded grey/blue region represents ± 1 sd, stimulus bar represents peristimulus duration, shaded green region represents our standard 50-150 analysis window used in previous figures). B) The varying facilitation onset time courses for 40 repeated trials in a single CSTMD1 neuron (grey lines) and the mean (black line). Red vertical dashed line represents short path onset. C) The mean facilitation onset time courses in CSTMD1 across 9 dragonflies. D) Neuronal response in the standard analysis window for each trial for long and short trajectories over the duration of a long experiment in a single CSTMD1 recording. E) Response variance as a function of mean spike count for the peristimulus duration of a long path trial, calculated by a 100 ms bin slid at 1 ms increments. Dot color represents the center time point of the window (Red t = 0s, Green t = 1s). F) Fano factor calculated for both short and long paths, calculated with 100 ms bins. Vertical Red dashed line represents the short path onset. G) Fano factor for the short path trial (same data as F) superimposed on the normalized response onset (same data as B), showing the inverse relationship between the strength of gain modulation and the magnitude of neuronal variability.

All statistical tests are non-parametric, paired, two-tailed and corrected for multiple comparisons.

## Results

### Spatiotemporal tuning of gain modulation

To investigate which target properties affect the magnitude of gain modulation, we present target trajectories segmented into two sections; a primer and a probe (Figure 1A-C). Responses to ‘probe’ targets are weak when presented alone, and enhanced when preceded by a ‘primer’ of varying durations or distances (Figure 1D).

Previous studies of STMD facilitation presented priming targets that drifted for similar durations and covered similar distances (Nordström et al. 2011; Dunbier et al. 2012; Wiederman, Fabian et al. 2017). These experiments demonstrated consistent increases in local gain, however confound the primer duration with the space it traversed. Therefore, it is unclear whether gain modulation results from a target stimulating a certain number of detectors across space, or whether it simply requires constant stimulation over a sufficient duration of time. To answer this question, we present primer targets that drift for different combinations of space and time, each terminating at the start position of a probe stimulus. By quantifying the change in probe response elicited by each primer we observed the strength of gain modulation (figure 2A).

If gain modulation only requires the stimulation of several adjacent motion detectors (~5° and larger), we should observe strong facilitation irrespective of the target’s duration traversing that space. Conversely, if gain is modulated only by the time of stimulation, primers of equal duration (over varying distance) should produce equal facilitation. As the primer’s duration or span approach zero, we know that no facilitation will occur (i.e. the probe alone response). We therefore limited the range of test durations and spans to elicit responses from an STMD neuron. Our results reveal that all of the tested primers produce an increase in gain (figure 2B). Pooling trials where the primer covered an equal length or duration reveals no net effect on the amount of facilitation generated. However, if we sort trials by primer velocity (primer length divided by primer duration) we observe responses similar to a velocity tuning function. Primers that drift at 80°/s produced the strongest increase in gain, irrespective of their duration or length. This velocity dependence suggests a motion-input pathway underlying the facilitation, on a scale less than 5° and elicited in under 125 ms.

### Target velocity and the spread of gain

A distinguishing property of predictive gain modulation is that when a moving target is temporarily occluded, the ‘focus’ of gain modulation shifts forward in space (Wiederman, Fabian et al. 2017). This extent of the forward spread matched the distance the target would have travelled had it continued at the same velocity during the two tested occlusion periods (150 ms and 300 ms). However, because this experiment was conducted at a single primer velocity, it is unclear whether the focus spreads at a target-matched velocity, or a constant ‘hardwired’ velocity that by chance matched our primer velocity. To investigate further, we present two primers of equal duration, one drifting at 30°/s and the other at 60°/s (Figure 3A). The faster primer covers twice the distance of the slow primer, however terminates at the same position. Following termination of the primer, we introduce a pause of either 100 ms or 300 ms to allow gain to spread, before presenting probes at different locations ahead and behind the primers final position (Figure 3B). If the predictive focus spreads at a primer matched velocity, then over the same time period the gain elicited by the faster primer should spread twice as far as the gain elicited by the slower primer.

Similar to the results presented in figure 2, the primer velocity has a large effect on the size of the gain modulation (Figure 3C). As the 60°/s primer lies closer to the velocity optimum, it evokes stronger faciliatory effects across all positions tested. This results in a vertical offset in the response change for the faster primer, potentially masking shifts in space that are due to the velocity of the stimulus. We corrected for the different strength primers by subtracting the mean difference between primer strength from the 60°/s primer trials at all positions following the 100 ms (Figure 3D) and 300 ms (figure 3E) pauses. The gain is modulated (change in response) by either the 30°/s (green line) or 60°/s (red line) primer and after either a 100 ms (Figure 3C) or 300 ms (Figure 3D) pause. As previously reported, a longer pause results in a spread forward of the gain modulation (Wiederman, Fabian et al., 2017). However, we did not observe a statistically significant effect of the primer velocity on the distance of this predictive spread. The spatial distribution of gain modulation is similar across both velocities tested with only the consistent upwards offset across all jump sizes.

### Gain modulation in response to the height of the primer target

Dragonflies are highly selective for the angular size of prey, with pursuit rarely initiated for targets spanning >1° on the retina (Combes et al. 2013; Lin and Leonardo 2017). The tuning range of STMD neurons is usually much broader, with CSTMD1 producing optimal responses to targets spanning 2-3° (Geurten et al. 2007). In the past, target height tuning has been measured with targets that drift across the entire visual display and are therefore ‘self-facilitated’. This means that these tuning curves represent not only the underlying size tuning of CSTMD1, but also any potential size tuning of gain modulation itself. During pursuit of prey or conspecifics the size of a target will fluctuate, gradually increasing in angular size as the distance to the target is reduced. Given that predictive gain modulation shifts CSTMD1’s direction tuning to match the direction of the target being tracked (Wiederman, Fabian et al. 2017), it is possible that a targets angular size could also shift the optimal target height? Such an effect would be similar to that observed in feature based attention studied in the human visual cortex (Saenz et al, 2002).

Here we investigated whether gain modulation produced by primers of different height affects CSTMD1’s height tuning. We present a series of probes of varying heights, either alone or following the presentation of primers of two heights, either 0.5° or 5° (Figure 4A,B). These sizes lay either side of CSTMD1’s optimal height tuning, and are expected to elicit responses of similar magnitude (Geurten et al. 2007).

Our results show that CSTMD1’s optimum target height remains at 2-3°, irrespective of whether or not the target is primed, or the size of the primer itself (Figure 4C). Both primers produced responses of equal strength (Figure 4d, 123.4 ± 16.9 for 0.5°, 123.7 ± 15.6 for 5°, P = 0.84). However, the strength of the response gain was not equal across all probe sizes. Rather, the gain produced by a given primer is greatest when the probe is at an optimal size and weaker when probes are either too small or too large (39.6±6.8 for weak probe sizes, 84.8± 13.5 for strong, p=0.007).

### Gain modulation and neuronal variability

Following our description of the effect of primer attributes (e.g. length, duration, velocity and height) on mean predictive gain modulation, we turned to an investigation of the response variance (both inter-trial and inter-animal). That is, what variability is observed in this response gain, when such primer parameters are kept constant? Nordström et al. (2011) reported that gain in CSTMD1 builds with a mean time to 50% of maximum response (t-50) of 180 ms, and this value justifies our time window for analysis in all subsequent work. However, the question arises as to whether this value is consistent across multiple animals or stable within the same neuron over extended periods of time.

We measured variability in the onset time course of gain modulation, with the presentation of a series of targets drifting on either short or long paths (figure 5A). We can determine the time onset of gain modulation, by normalising the short path responses by their corresponding long path counterparts (the spatial location of the short path covers the long path’s second half). This approach provides a temporal signature (onset time course), whilst accounting for inhomogeneity of the spatial receptive field. We repeated this experiment 40 times in a single CSTMD1 recording at regular intervals over 90 minutes (figure 5A,B), and across 9 CSTMD1 recordings in 9 different animals (5C). In both cases we observe a variability in the response time course, even though the underlying stimuli and neuronal architecture are identical. This raises the intriguing possibility that the strength of the predictive gain modulation is itself modulated by an internal stimulus-independent factor. Variations in the response onset time course will result in variance in the number of spikes in our standard analysis time window (50-150 ms following stimulus onset). A degree of STMD variability may be due to neuronal habituation, resulting from the presentation of repeated stimuli. Even though each stimulus was separated by a 146 s rest period, responses show a slight downward trend due to some habituation (Figure 5D). Additionally there is variation in response over time, especially for short path targets (black line). To quantify the variability of responses over time we plot the response variance as a function of mean spike count for the repeated presentation of a target trajectory in a single CSTMD1 neuron (Fig 5E, same data as figure 5A). This data is generated by sliding a short analysis window across the 1s peristimulus time window in 1 ms increments. The relationship between response variance and mean spike-count varies dependent on the length of analysis window (Warzchecha and Egelhaaf, 1999), therefore we analyse our data across a broad range of bin sizes (100 ms bin data shown). We see that the ratio between variance and mean spike-count is strongly dependent on the time following stimulus onset, with very large values (Fano Factor > 4) shortly following stimulus onset, dropping to < 0.5 after several hundred milliseconds of motion (Figure 5F). The time course of this Fano factor drop is closely matched to the build-up of gain modulation following the onset of both the short and the long paths (Figure 5G). Given that both paths cover the exact same space on the retina this difference in neuronal variability cannot be explained by input noise. Instead, this variation arises from changes in the strength of modulatory signals throughout the visual pathway.

## Discussion

The responses of individual STMD neurons are generated by complex interactions between the parameters of a stimulus and the spatiotemporal dynamics of the receptive field (Wiederman, Fabian et al. 2017). Our previous description of STMD gain modulation proposed that neuronal responses increased over hundreds of milliseconds due to the continuous target motion (Nordström et al. 2011). Here we show that this increase is not a simple time-dependent gain modulation, instead displaying dependencies on priming target velocity.

We also note that we have observed the same primers generating different amounts of facilitation across probes with different parameters, such as angular size. This result suggests that gain modulation is not a simple additive operation. Instead, the magnitude of gain modulation appears to be dependent on both the salience of a priming stimulus and the salience of the probe that follows. This is in agreement with our previous work, where the same priming stimulus generated different strengths of gain modulation for probes of different contrasts and directions (Wiederman, Fabian et al, 2017). In addition, when all conditions are kept constant we see that the magnitude and time course of response onset varies considerably. Since targets moving on long and short paths stimulate identical presynaptic visual pathways this large increase in variability must be the result of variability in internal modulatory signals over time. This finding is consistent with prior work that concluded that most response variability in higher-order processing areas is generated by variability of modulatory signals rather than input noise (Goris et al., 2014). At this time the neuronal circuits responsible for modulating this gain magnitude are unknown. Target tracking and pursuit simulations which implement a dragonfly inspired predictive gain modulation mechanism found that the optimal facilitation time constant was variable, depending on the spatial statistics of a scene (Bagheri et al., 2015). In cluttered scenes a longer time constant increases performance due to the high likelihood of temporary target occlusions, however in more sparse scenery a rapid facilitation time course is beneficial, as the target remains highly discriminable. While the spatial statistics of our visual stimulus are identical across each trial these simulations suggest that the ability to dynamically tune the time course of gain modulation via an unknown pathway could be a useful tool for target detecting systems.

The downside of a dynamically modulated gain mechanism is that neuronal signals to repeated stimuli are significantly more variable. In most sensory neurons variance increases with mean spike count, with a slope that varies greatly depending on many conditions (Aldo Faisal et al., 2008). In H1 of the fly, the ratio between variance and mean spike count (also known as Fano factor) has been reported as approximately 0.16 for 10 ms analysis windows, and 0.07 for 100 ms windows, suggesting that repeated stimuli are encoded with relatively low variability (Warzecha and Egelhaaf 1999). CSTMD1’s target responses are extremely unreliable until several hundred milliseconds have passed, imposing an interesting challenge to potential target tracking control systems. However while the changing magnitude of this gain mechanism may increase neuronal variability in an early time window, it may be significantly decreasing neuronal variability in late time windows, as responses are driven closer to saturating regions of CSTMD1s firing range.

The key advantages of such a non-linear stimulus-stimulus interaction is that robust tracking of targets can be obtained amidst dynamically changing conditions. A target that temporarily drops out of an optimal range in size, contrast or velocity will still elicit strong responses due to enhanced local gain generated during its prior path. Conversely, in contrast to our previous ideas, a target that abruptly enters the visual field can be detected in a reasonably short time if it accurately matches the systems tuning parameters. Such a stimulus-stimulus interaction is favourable for improving signal-to-noise ratio when extracting small moving targets from cluttered backgrounds.

